# Synthetic essentiality of TRAIL/TNFSF10 in VHL-deficient renal cell carcinoma

**DOI:** 10.1101/2025.05.29.621197

**Authors:** Xuechun Wang, Loan Duong, Yujing Qin, Rossella Parrotta, Paresh Kumar Purohit, Yihao Fang, Guoqiang Liu, Jianping He, Jiling Wen, Yan Liu, Yuting Zhang, Junling Zhao, Zachary T Schafer, Xuemin Lu, Eva Szegezdi, Xin Lu

## Abstract

Clear cell renal cell carcinoma (ccRCC) is the most common and aggressive subtype of kidney cancer. Loss of *von Hippel–Lindau* (*VHL)* and the consequent activation of hypoxia-inducible factor-α (HIFα, especially HIF2α) plays an essential role in ccRCC initiation and progression. The approved HIF2α inhibitor belzutifan faces the challenge of resistance, presenting an opportunity of co-targeting HIF2α and another vulnerability. This study elucidates the synthetic essentiality of TRAIL (tumor necrosis factor-related apoptosis-inducing ligand) in VHL-deficient ccRCC, uncovering a novel reciprocal regulation between HIF2α and TRAIL. TRAIL was identified as a direct transcriptional target of HIF2α and paradoxically found to be crucial for cell proliferation, primarily by activating the p38 MAPK pathway and facilitating G1/S phase transition. Depletion of endogenous TRAIL or inhibition of HIF2α with belzutifan sensitizes ccRCC cells to recombinant TRAIL, presenting a promising avenue for combination therapy to overcome both TRAIL resistance and belzutifan resistance in treating ccRCC.

**Graphical Abstract:** 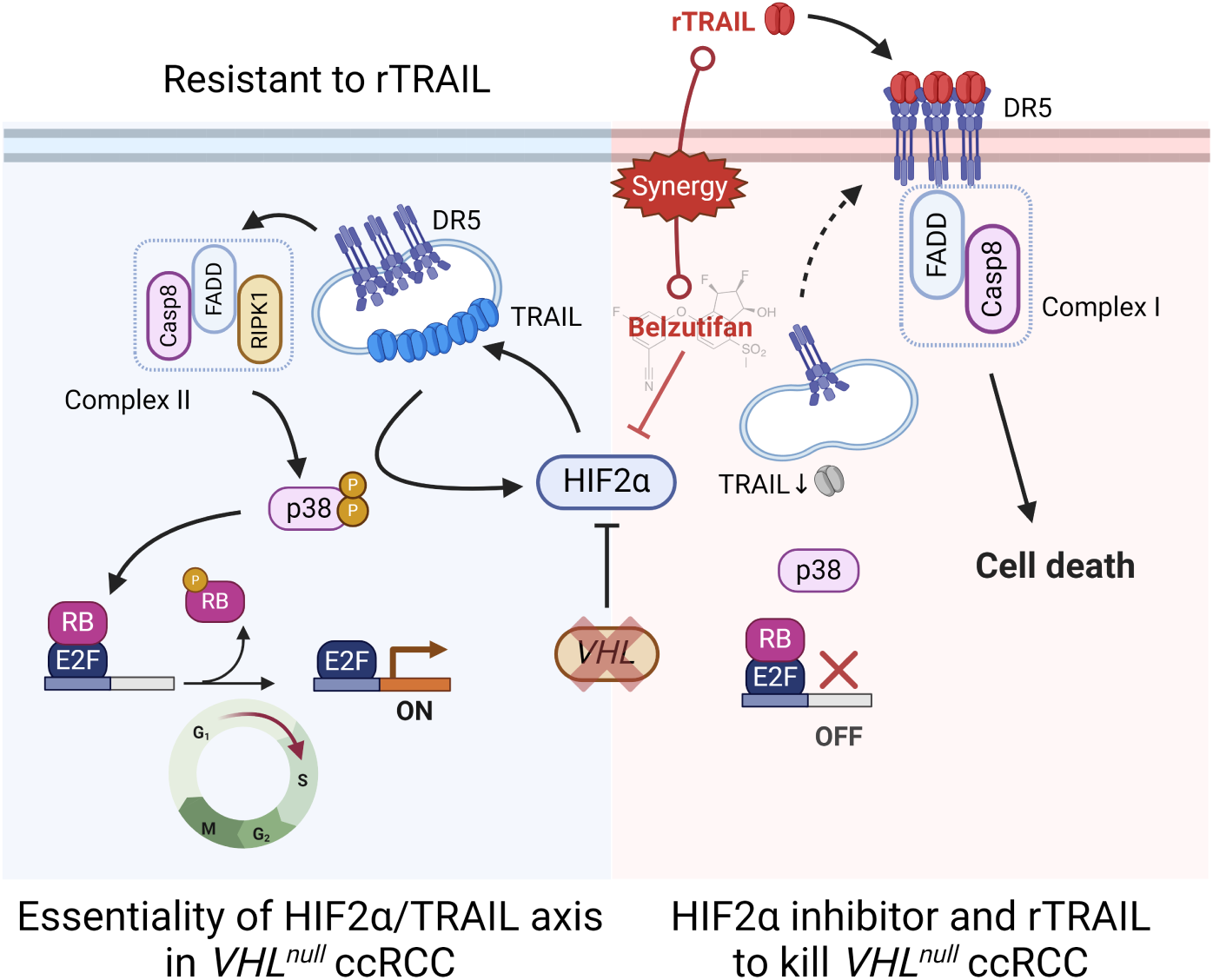

## Introduction

Kidney cancer, also known as renal cell carcinoma (RCC), represents ∼2% of all cancer diagnoses and cancer deaths globally, affecting over 434,000 individuals worldwide and causes over 155,000 deaths annually ^1^. Clear cell renal cell carcinoma (ccRCC) is the most common and aggressive subtype of RCC, accounting for 75-80% of RCC diagnoses ^2^. ccRCC occurs in both sporadic and familial forms. Patients affected by von Hippel–Lindau (VHL) disease are at high risk of developing multifocal, bilateral ccRCC ^3^. Loss or mutation of the tumor suppressor gene *VHL* is an obligate initiating step for ccRCC ^2,4^. *VHL* inactivation leads to stabilization of hypoxia-inducible factor-α (HIFα, primarily HIF1α and HIF2α) and a few other recently identified targets ^5–7^. HIF transcription factors activate many oncogenic targets promoting malignancy traits such as hyper-proliferation, angiogenesis, Warburg-like metabolic reprogramming, and metastasis ^2,4,8^. In ccRCC, HIF2α appears to play a more dominant tumor-promoting role than HIF1α, prompting the development of the HIF2α inhibitor belzutifan which was approved to treat patients with VHL disease tumors ^9–11^. Despite the approval, resistance to belzutifan by ccRCC remains a challenge and presents an urgent need to identify other vulnerabilities in ccRCC that may be co-targeted with HIF2α inhibition ^9–11^.

TRAIL (tumor necrosis factor-related apoptosis-inducing ligand), encoded by gene *TNFSF10*, is a death ligand cytokine and a promising anti-cancer agent due to its ability to selectively kill cancer cells without impacting normal cells^12^. Human TRAIL binds to four membrane receptors TRAIL-R1 (DR4), TRAIL-R2 (DR5), TRAIL-R3 (DcR1), and TRAIL-R4 (DcR2), as well as one soluble receptor, osteoprotegerin (OPG) ^13^. Only DR4 and DR5 have full-length intracellular death domains (DD) and mediate TRAIL-induced apoptosis, while the other receptors act as TRAIL decoys to lower TRAIL availability for DR4 and DR5. In the canonical TRAIL-induced apoptosis pathway, TRAIL binds to the ligand binding extracellular domain of DR4 or DR5 and initiates receptor oligomerization, which triggers apoptotic signaling cascade by recruiting FAS-associated death domain (FADD) and pro-caspase-8 to form the death-inducing signaling complex (DISC). The TRAIL DISC induces the activation of pro-caspase-8, thereby activating the caspase cascade. Induction of apoptosis can be inhibited by FLICE-like inhibitory protein (cFLIP) because it competes with pro-caspase-8 for FADD. Conversely, TRAIL can also activate a non-canonical pathway in some cells and contribute to proliferation/pro-survival signals ^14^. In the non-canonical pathway, TRAIL triggers the formation of a secondary complex which retains the DISC components FADD and pro-caspase-8 but recruits several other factors, including receptor-interacting serine/threonine protein kinase 1 (RIPK1), TRAF2, and NEMO/IKKγ, which in turn activate kinases such as IKK, JNK, and p38 mitogen-activated protein kinase (MAPK) ^15^. Due to the disparate effects of the canonical and non-canonical TRAIL pathways, resistance to TRAIL treatment is common in cancers including ccRCC ^16^, and clinical trials testing TRAIL-based therapeutics showed limited efficacy ^17^. A potential strategy is the rational sensitization of cancer cells to TRAIL through combination therapy.

Synthetic essentiality is a phenomenon in cancer where a gene is non-essential under normal conditions but becomes essential for cell survival or proliferation in the presence of a specific genetic alteration ^18^. It has been reported in PTEN-deficient prostate cancer^19^ and rhabdomyosarcoma^20^, and APC-deficient colorectal cancer^21^, which reveal novel cancer vulnerabilities and therapeutic opportunities. For ccRCC, while targets synthetic lethal with VHL loss are reported (CDK4/6, TBK1) ^22,23^, synthetic essentiality genes for VHL-deficient ccRCC have not been explicitly defined. In this study, we were inspired by the paradoxical essentiality of TRAIL in VHL-null ccRCC cell lines, and identified the reciprocal positive regulation of HIF2α and TRAIL. TRAIL promotes ccRCC cell proliferation through activating p38 MAPK and G1/S transition. Critically, abrogating endogenous TRAIL in ccRCC cells, through direct TRAIL knockdown or HIF2α inhibition, sensitizes ccRCC cells to recombinant TRAIL treatment, paving a translational path of combination therapy in ccRCC.

## Results

### TRAIL is a HIF2α target and a candidate synthetic essential gene

To identify putative synthetic essential genes in VHL-deficient ccRCC, we focused on 786O cells (the most commonly used VHL-deficient ccRCC cell line) and generated a HIF2α (encoded by *EPAS1*) knockout subline (via CRISPR/Cas9) and determined the gene expression changes induced by HIF2α deficiency by RNA-seq. Compared with 786O-sgEPAS1, 786O-Ctrl enriched various molecular pathways, including E2F targets, G2M checkpoint, hypoxia, and glycolysis, which were also enriched when 786O-Ctrl was compared with VHL-restored 786O subline as we previously reported ^7^ (**Supplementary Fig.1A, Supplementary Table 1**), confirming the validity of HIF2α knockout. We prioritized candidates for synthetic essential genes based on two criteria: (i) significantly downregulated by HIF2α knockout compared with 786O control cells (i.e., HIF2α-upregulated genes); (ii) being a preferentially essential gene in 786O cells based on Dependency Map (DepMap) ^24^. The intersection of the two criteria yielded 57 potential synthetic essential genes (**Fig. 1A, Supplementary Table 2**), with TNFSF10/TRAIL being the top candidate (**Fig. 1B, Supplementary Fig. 1B**).

**Figure 1.**
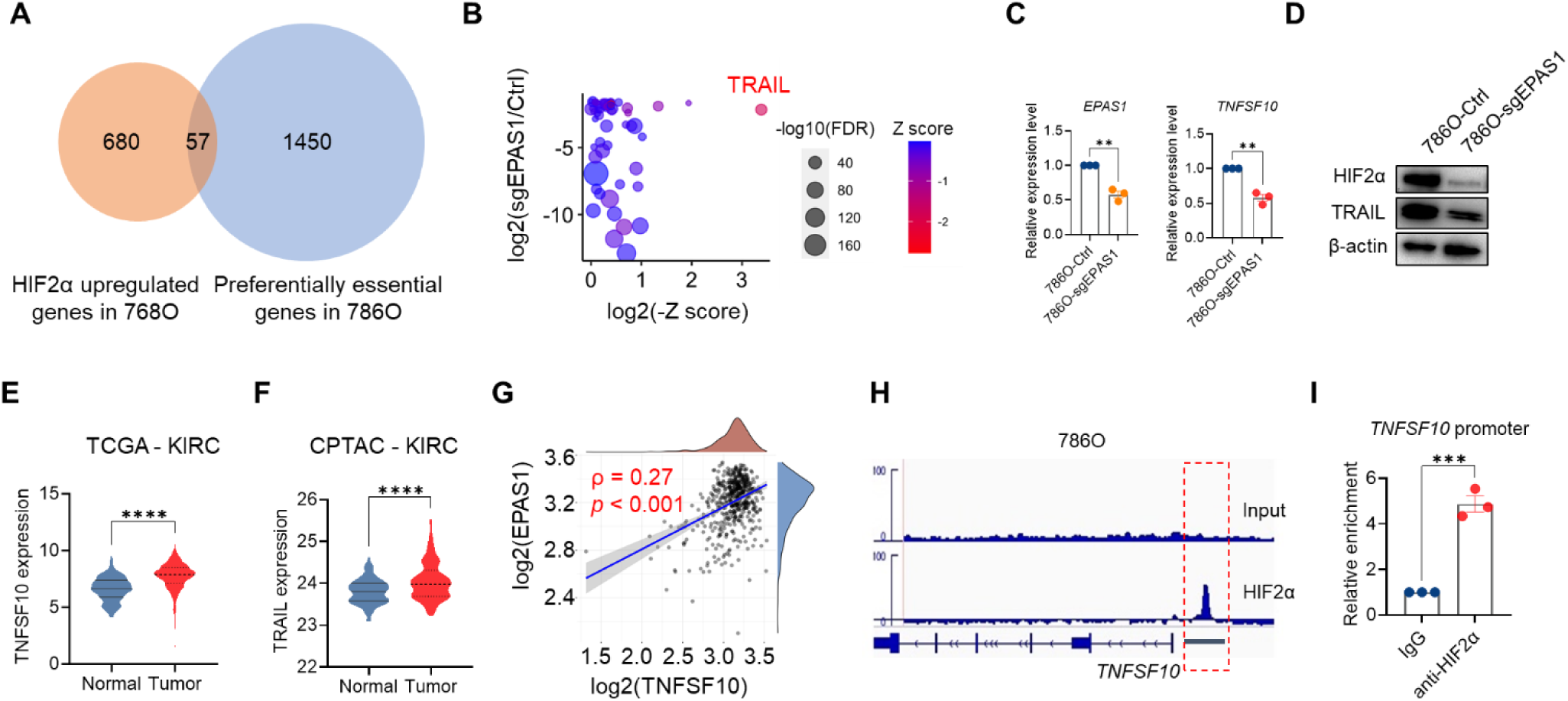
TRAIL is a HIF2α target and a candidate synthetic essential gene. **A.** Venn diagram between HIF2α-upregulated genes (logFC(Ctrl/sgEPAS1) > 1.5, FDR < 0.05) and DepMap preferentially essential genes (gene effect < 0, Z-score < -1) in 786O. **B.** Bubble plot of the 57 overlapped genes. The bubbles’ size and color reflect the false discovery rate (FDR) and Z-score, respectively. TRAIL has lowest Z-score and is highlighted. **C.** Relative mRNA expression of *EPAS1* and *TNFSF10* in control and HIF2α-knockout 786O sublines (n=3), measured with qRT-PCR. **D.** Immunoblotting of HIF2α and TRAIL in control and HIF2α-knockout 786O sublines. **E.** Relative mRNA expression of *TNFSF10* in normal (n=161) and tumor tissues (n=532) of TCGA KIRC patient’s cohort. **F.** TRAIL protein expression in normal (n=84) and tumor tissues (n=110) of CPTAC KIRC patient’s cohort. **G.** Scatter plot of *TNFSF10* and *EPAS1* levels in the TCGA KIRC cohort. Spearman correlation coefficient ρ and P value are marked. **H.** HIF2α and input ChIP-Seq signals in the *TNFSF10* locus with the promoter region chr3:172,242,513 - 172,243,623 highlighted in the red box. **I.** ChIP-qPCR result for IgG and anti-HIF2α at the *TNFSF10* promoter (n=3). In **C** and **I**, data represent mean ± SEM. Unpaired t-test for **C** and **I**, Mann-Whitney test for **E** and **F**, **P<0.01, ***P<0.001, ****P<0.0001.

The downregulation of TRAIL by HIF2α knockout was confirmed at the mRNA (**Fig. 1C**) and protein levels (**Fig.1D**). TRAIL expression was also examined in primary kidney renal clear cell carcinoma (KIRC) RNA-seq data of The Cancer Genome Atlas (TCGA) ^25^ and the proteomics data of Clinical Proteomic Tumor Analysis Consortium (CPTAC) ^26^. There was a significant increase in TRAIL expression in ccRCC compared with normal kidney tissue at both mRNA (**Fig. 1E**) and protein levels (**Fig. 1F**), suggesting a potential pro-tumor role of TRAIL in renal carcinogenesis. Moreover, there was a positive correlation between *TNFSF10* and *EPAS1* expression in TCGA KIRC data (**Fig. 1G**), supporting the regulation of TRAIL by HIF2α. To examine whether TNFSF10 is a direct transcriptional target of HIF2α, we re-analyzed the HIF2α ChIP-seq data generated in 786O cells ^27^ and identified the association of HIF2α with the promoter region of *TNFSF10* (**Fig. 1H**). The binding of HIF2α to the promoter regions of *TNFSF10* and *GLUT1* (a known HIF2α target) was validated with ChIP-qPCR (**Fig.1I**, **Supplementary Fig.1C**). Together, TRAIL is a direct transcriptional target of HIF2α and a top-ranked candidate synthetic essential gene.

### TRAIL depletion attenuates ccRCC proliferation *in vitro* and tumor growth *in vivo*

To assess TRAIL essentiality in VHL-null ccRCC cells, we started with constitutive TRAIL knockdown with the short hairpin RNA (shRNA) approach. Each of the three shRNAs (#24, #25, #27) targeting different TRAIL transcript regions reduced TRAIL levels effectively (**Supplementary Fig. 2A-B**) and dramatically lowered colony formation ability (**Supplementary Fig. 2C**). Consequently, no 786O sublines with sustained TRAIL knockdown could be obtained. Therefore, we generated a doxycycline-inducible shTRAIL (dishTRAIL) subline of 786O using one of the shRNA sequences (#25) and confirmed the inducibility and efficacy of the knockdown (**Fig. 2A-B**). While doxycycline did not affect unmodified 786O cells (**Supplementary Fig. 2D**), doxycycline-induced TRAIL knockdown in 786O-dishTRAIL cells slowed cell proliferation (**Fig. 2C**) and diminished colony formation potential (**Fig. 2D, Supplementary Fig. 2E**). Consistent with the *in vitro* effect, orthotopic tumor formation by 786O-dishTRAIL cells (labeled with a luciferase report gene) was impeded in mice fed with doxycycline diet compared with control diet (**Fig. 2E-F**). These results revealed the paradoxical role of TRAIL in 786O and prompted us to explore the mechanism.

**Figure 2.**
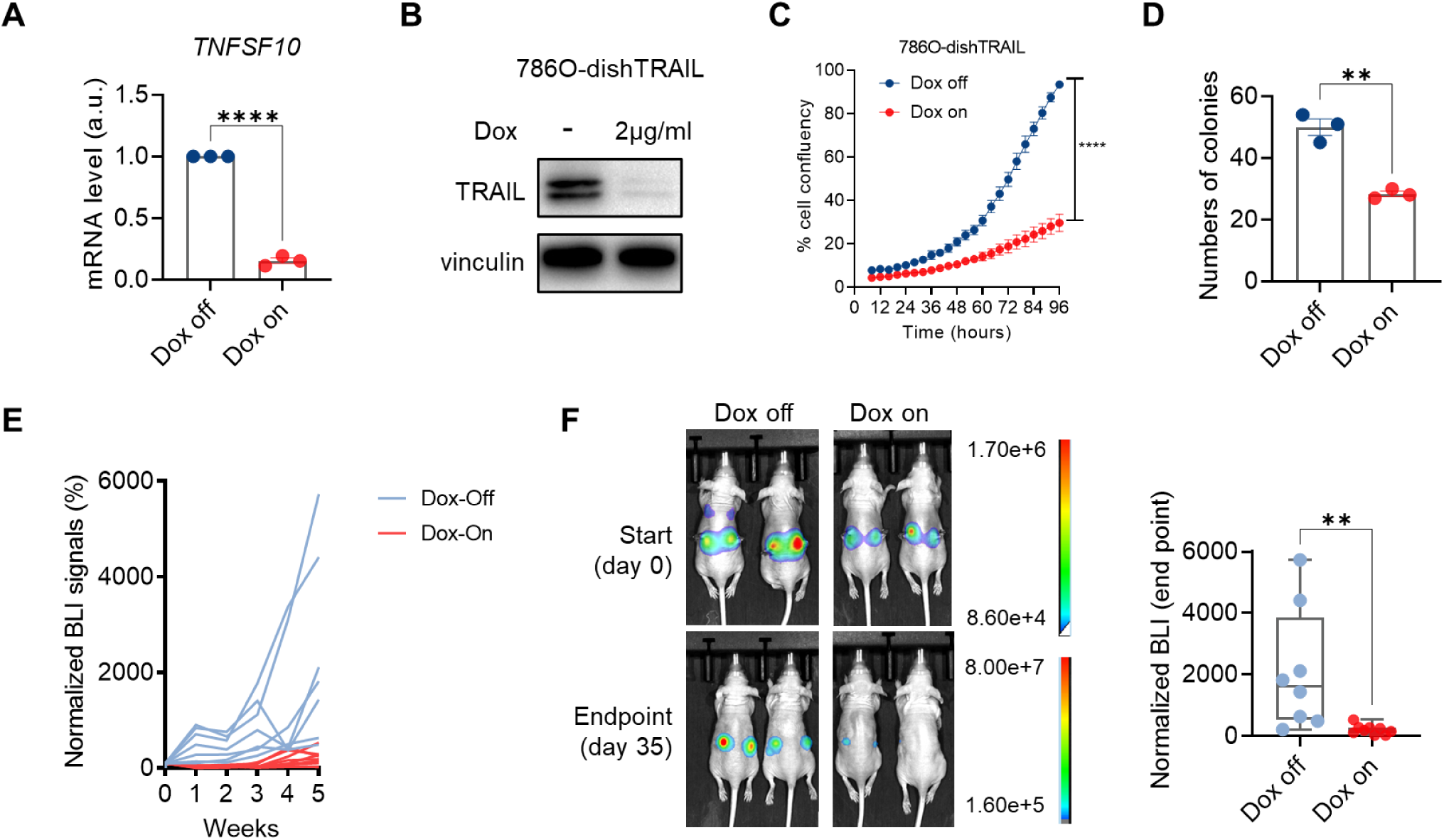
TRAIL depletion attenuates ccRCC proliferation in vitro and tumor growth in vivo. **A.** qRT-PCR result showing relative mRNA expression of *TNFSF10* of 786O-dishTRAIL cells under doxycycline off or on (2 μg/ml, 24 hours) conditions (n=3). **B.** Immunoblotting of TRAIL of 786O-dishTRAIL cells under doxycycline off or on (2 μg/ml, 48 hours) conditions. **C.** Cell proliferation curves of 786O-dishTRAIL cells under doxycycline off or on (2 μg/ml) conditions over a 96-hour period. Each data point was calculated from the cell confluency images (n=6). **D.** Clonogenic assay showing numbers of colonies (diameter of colony > 0.2cm) of 786O-dishTRAIL cells under doxycycline off or on (2 μg/ml, 10 days) conditions (n=3). **E.** Spider plots showing normalized bioluminescence imaging (BLI) signals for nude mice orthotopically injected with 786Odish-TRAIL cells and fed with doxycycline diet (Dox on, n=5) and control diet (Dox off, n=4). **F.** Representative BLI images (day 0 and 35) and normalized BLI signals (day 35) for Dox off and Dox on groups. In **A, C, D** and **F**, error bars represent SEM. Unpaired t-test for **A, C** and **D**, Mann-Whitney test for **F**, **P<0.01, ****P<0.0001.

### TRAIL activates p38 MAPK and facilitates G1/S progression in ccRCC cells

To identify the mechanism underlying the preferentially essential role of TRAIL in ccRCC, we conducted transcriptomic profiling of 786O-dishTRAIL cells with or without doxycycline treatment (48 hours). RNA-seq identified 484 differentially expressed genes after TRAIL knockdown (|Fold Change| > 1.5 and FDR < 0.05), including 282 upregulated and 202 downregulated genes (**Fig. 3A, Supplemental Table 3**). Gene Set Enrichment Analysis (GSEA) using the MSigDB Hallmark gene sets for the differentially expressed genes identified MYC targets and E2F targets as the most significantly enriched gene sets for Dox-off TRAIL^high^ condition (**Fig. 3B, Supplementary Fig. 3A, Supplemental Table 4**). MYC is a master regulator of cancer hallmarks ^28^, whereas E2F functions as the core transcriptional machinery driving cell cycle progression ^29^. Both MYC and E2F are the key effectors downstream of HIF2α in ccRCC ^30^. The classical function of TRAIL is to induce apoptosis. However, inducible knockdown of TRAIL in 786O did not induce apoptosis (**Fig. 3C**), whereas staurosporine did (**Supplementary Fig. 3B**). This result suggests that TRAIL regulates ccRCC cells through non-apoptotic mechanisms.

**Figure 3.**
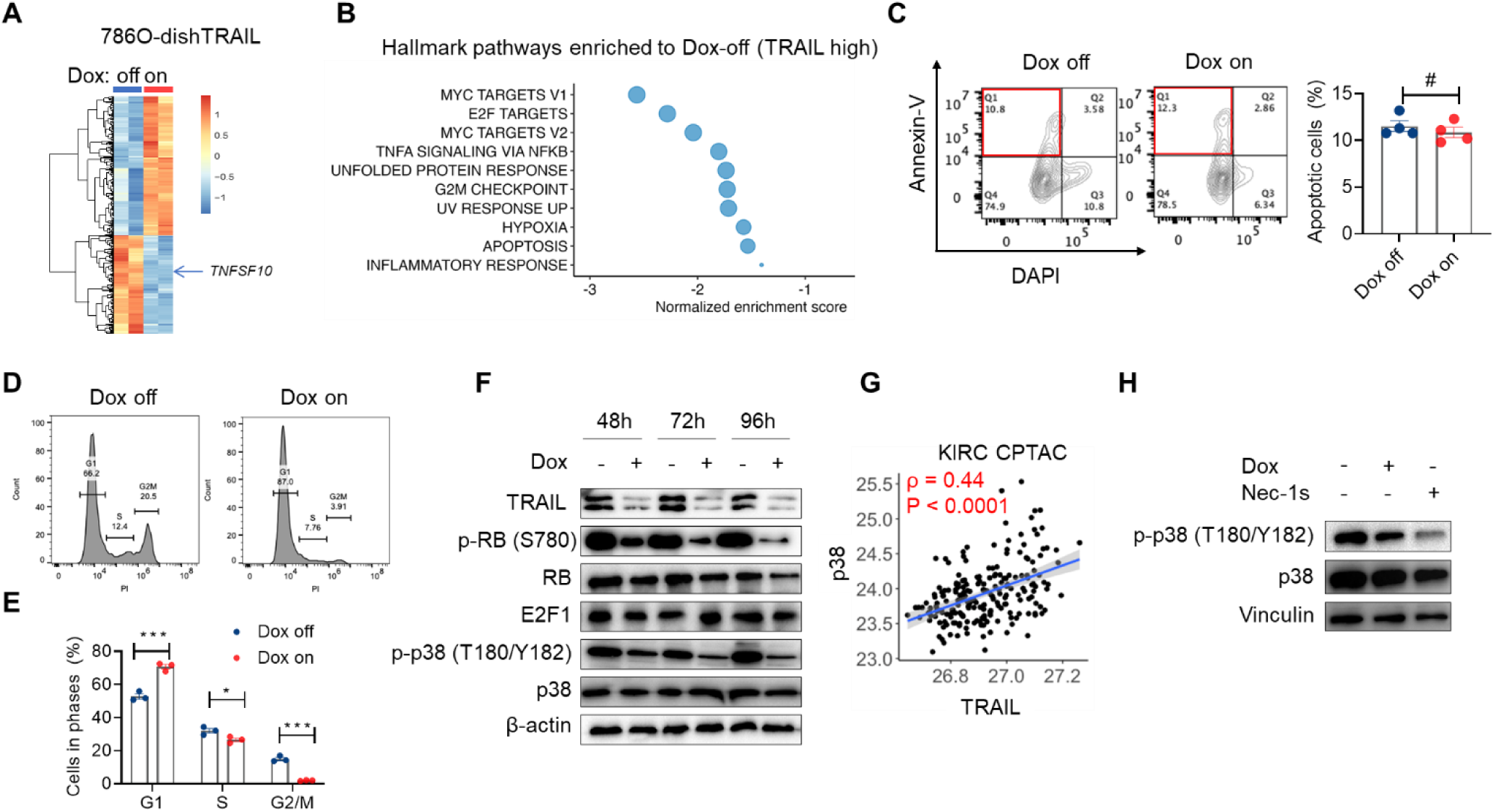
TRAIL activates p38 MAPK and facilitates G1/S progression in ccRCC cells. **A.** Heatmap showing differential gene expression (|fold Change| > 1.5 and FDR < 0.05) between 786O-dishTRAIL treated with doxycycline (Dox on, 48 hours) and untreated controls (Dox off). *TNFSF10* is among the downregulated genes by TRAIL knockdown. **B.** GSEA result showing MSigDB hallmark pathways (P < 0.05, FDR < 0.25) enriched to Dox-off condition. **C.** Representative flow cytometric plots of annexin-V-APC and DAPI staining on 786O-dishTRAIL under Dox on (48 hours) and Dox off conditions (n=4) and qualification of the percentage of apoptotic cells (red box). **D-E.** Representative flow cytometric plots for DNA content analysis using propidium iodide (PI) staining of 786O-dishTRAIL under Dox on (48 hours) and Dox off conditions (n=3), based on which the proportions of three cell cycle phases were quantitated. **F.** Immunoblotting showing changes of the indicated proteins and phosphorylations for 786O-dishTRAIL under Dox on and Dox off conditions for various durations (48, 72, 96 hours). **G.** Scatter plot of protein levels of TRAIL and p38 in ccRCC CPTAC data. Pearson correlation coefficient ρ and P value are indicated. **H.** Immunoblotting of p38 and phorpho-p38(Thr180/Tyr182) for 786O-dishTRAIL treated with 10µM necrostatin-1s (Nec-1s) for 1 hour or 48 hours of doxycycline. In **C** and **E**, error bars represent SEM. Unpaired t-test, *P < 0.05, ***P < 0.001, ^#^P > 0.05.

We investigated the effect of TRAIL on cell cycle regulation with flow cytometric DNA content analysis (**Fig. 3D**). Doxycycline-induced TRAIL knockdown increased the proportion of cells in G1 phase and decreased the proportions in S and G2/M phases (**Fig. 3E**), indicating that TRAIL-depleted cells were arrested in G1/S transition. Consistent with the cell cycle phenotype, inducible TRAIL knockdown lowered cyclin D1 and cyclin E1 levels (**Supplementary Fig. 3C**). The retinoblastoma tumor suppressor protein (RB) controls the G1/S transition and inhibits cell cycle entry. RB phosphorylation by cyclin D-CDK4/6 and cyclin E-CDK2 inactivates RB and releases E2Fs to stimulate gene transcription and propel cell cycle^31^. Concordant with lower cyclin D1 and E1 levels, TRAIL knockdown decreased the RB phosphorylation without altering total RB or E2F1 levels (**Fig. 3F**). Therefore, TRAIL is essential for promoting G1/S transition and cell cycle through regulating the cyclin/CDK/RB/E2F axis.

To understand how TRAIL regulates the RB/E2F axis, we examined several downstream players of the non-canonical (non-apoptotic)TRAIL–TRAIL-R signaling, including p38 MAPK, ERK MAPK, IκB and NFκB ^13^. While ERK, IκBα and NFκB did not show changes upon TRAIL repression (**Supplementary Fig. 3D**), TRAIL knockdown reduced p38 MAPK phosphorylation while total p38 remained unchanged (**Fig. 3F**). TRAIL-induced p38 activation can produce pro-survival signals in prostate and breast cancer cells ^32,33^. Existence of the TRAIL-p38 regulatory axis in ccRCC was supported by the strong correlation between p38 and TRAIL proteins in KIRC CPTAC data (**Fig. 3G**). In the non-canonical TRAIL signaling, TRAIL-induced p38 activation depends on RIPK1 ^15^. In line with this, RIPK1 inhibition with necrostatin-1s reduced p38 phosphorylation in 786O cells, similar to the effect from TRAIL knockdown (**Fig. 3H**). p38 exerts dual roles in regulating cell cycle ─ while its activation often causes cell cycle arrest^34^, under certain cellular contexts p38 activation drives RB phosphorylation and G1/S transition ^35–37^. Taken together, the TRAIL-p38-RB-E2F axis emerges as a predominant mechanism for the pro-tumor function of TRAIL in ccRCC.

### TRAIL is required to sustain HIF2α expression and activity in ccRCC cells

The results above establish TRAIL as a transcriptional target and functional executor of HIF2α in ccRCC cells. Interestingly, inducible TRAIL silencing in 786O-dishTRAIL reciprocally reduced HIF2α protein level progressively (**Fig. 4A**), but *EPAS1* mRNA level was unaffected (**Supplementary Fig. 4A**). Consistently, HIF2α targets, *VEGFA* and *SLC2A1* were also downregulated by both HIF2α knockout and TRAIL knockdown (**Fig. 4B**). This result suggests that TRAIL might play a role in a feedback loop that mediates HIF2α activity. TRAIL likely regulates HIF2α protein level through p38, as p38 was shown necessary for hypoxia-induced or arsenite-induced HIF1α upregulation ^38,39^. To confirm p38 is essential for HIF2α expression and activity in 786O, we treated the cells with the p38 kinase inhibitor, SB203580. Similar to the effect of TRAIL knockdown, p38 inhibition reduced HIF2α protein levels (**Fig. 4C, Supplementary Fig. 4B**) and expression of *VEGFA* and *SLC2A1* (**Fig. 4D**), but did not affect *EPAS1* mRNA level (**Fig. 4D**). Furthermore, the positive correlation between TRAIL expression and hypoxia gene signature score (MSigDB Hallmark gene sets) across various cancer cell lines in the Cancer Cell Line Encyclopedia supports the role of TRAIL in regulating HIF signaling (**Supplementary Fig. 4C**). These results suggest a reciprocal regulation model of HIF2α and TRAIL.

**Figure 4.**
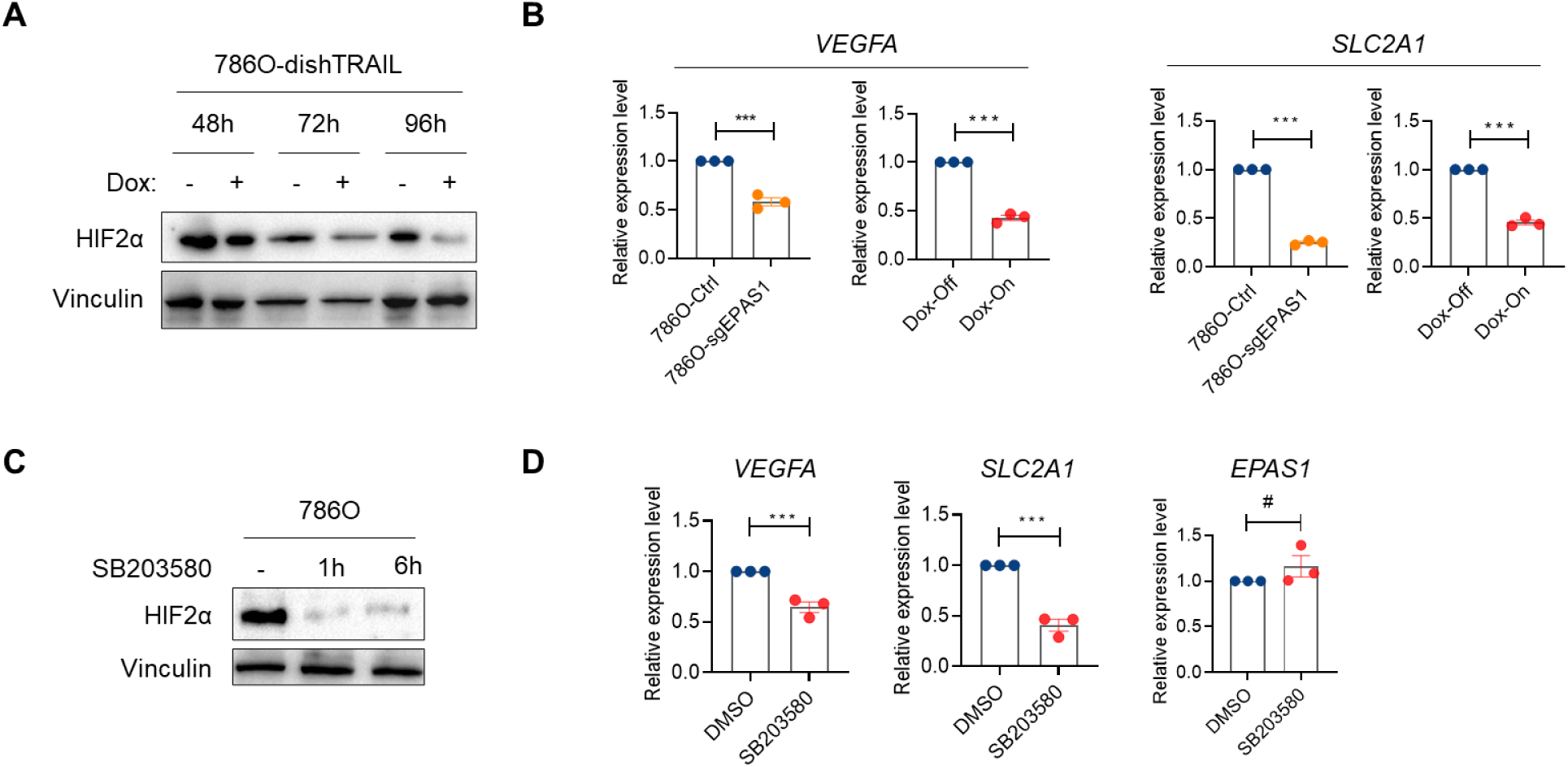
TRAIL is required to sustain HIF2α expression and activity in ccRCC cells. **A.** Immunoblotting of HIF2α for 786O-dishTRAIL treated with doxycycline (Dox on) and untreated controls (Dox off) across three time points (48,72, and 96 hours). **B.** qRT-PCR result of relative mRNA expression of *VEGFA* and *SLC2A1* for 786O-Ctrl, 786O-sgEPAS1, and 786O-dishTRAIL under Dox off and Dox on (96 hours) conditions (n=3). **C.** Immunoblotting of HIF2α for 786O treated with 25μM SB203580 for 1 hour or 6 hours. **D.** qRT-PCR result of relative mRNA expression of *VEGFA*, *SLC2A1* and *EPAS1* for 786O treated with 25μM SB203580 for 6 hours (n=3). In **B** and **D**, error bars represent SEM. Unpaired t-test, ***P < 0.001, ^#^P > 0.05.

### TRAIL depletion re-localizes DR5 to the plasma membrane

TRAIL induces cytoplasmic signaling through binding to either DR4 or DR5, among which DR5 has higher ligand binding affinity ^13^. Besides plasma membrane, DR5 can be found on the intracellular membranes, such as the endoplasmic reticulum (ER), endosomes and lysosomes, and non-membranous compartments such as nuclei; its subcellular localization may fine-tune the cell’s fate upon TRAIL binding ^40^. We found that high DR5 expression is associated with worse overall survival in ccRCC patients (**Supplementary Fig. 5A**). Both TRAIL and DR5 were localized abundantly in subcellular organelles of 786O cells and showed substantial colocalization (**Fig. 5A**). At least a sizable portion of TRAIL and DR5 were localized at lysosomes detected with the marker LAMP1 (**Fig. 5B-C**). Some DR5 was also localized in the ER as expected (**Supplementary Fig. 5B**). DR5 sequestration into intracellular localization is a strategy for cancer cells to evade TRAIL-induced apoptosis ^40^. After TRAIL knockdown in 786O cells, we observed an upregulation of DR5 on the plasma membrane (**Fig. 5D**), which was not the case for DR4 (**Supplementary Fig. 5C**). These results suggest that TRAIL contributes to the intracellular retention of DR5 and TRAIL depletion mobilizes DR5 to the cell surface.

**Figure 5.**
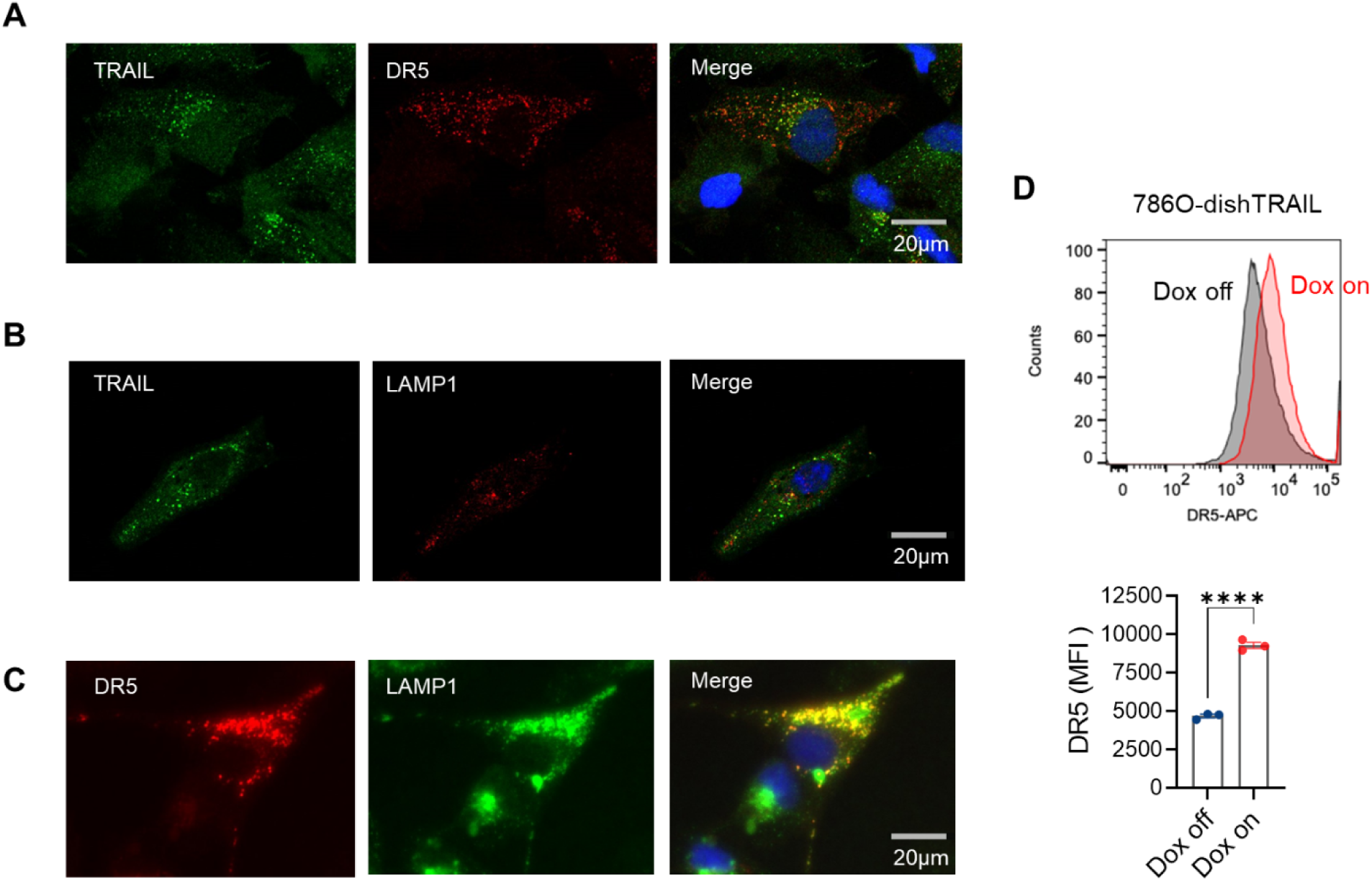
TRAIL depletion re-localizes DR5 to the plasma membrane. **A.** Immunofluorescence co-staining of TRAIL and DR5 in 786O. Scale bar 20μm. **B.** Immunofluorescence co-staining of TRAIL and lysosomal marker LAMP1 in 786O. Scale bar 20μm. **C.** Immunofluorescence co-staining of DR5 and LAMP1 in 786O. Scale bar 20μm. **D.** Representative flow cytometry plot and quantitation of mean fluorescence intensity (MFI) of DR5 signals on the cell surface in 786O-dishTRAIL cells under Dox on or Dox off conditions (48 hours, n=3). Error bars represent SEM. Unpaired t-test, ****P < 0.0001.

### HIF2α inhibitor synergizes with recombinant TRAIL to reduce 786O viability

The known resistance of 786O cells to recombinant TRAIL (rTRAIL)-induced apoptosis was at least partly due to low cell surface density of TRAIL receptors ^41^. The elevated DR5 expression on the surface of 786O cells upon TRAIL knockdown prompted us to test whether these cells became more sensitive to rTRAIL. Two different concentrations of rTRAIL were used to treat 786O-dishTRAIL in the absence or presence of prior Dox-induced TRAIL silencing. Loss of endogenous TRAIL expression sensitized the cells to killing by rTRAIL at 200ng/ml, which was more pronounced by 500ng/ml rTRAIL (**Fig. 6A**). Given that HIF2α upregulates TRAIL in 786O cells, we hypothesized that belzutifan (the FDA-approved HIF2α inhibitor) should reduce TRAIL expression and sensitize the cells to rTRAIL. Belzutifan reduced HIF2α expression in 786O cells as previously reported ^10^; but more importantly, belzutifan reduced TRAIL and phospho-p38 levels (**Fig. 6B**). While the viability of 786O cells was not affected by single treatment of rTRAIL or up to 2 mM belzutifan alone, combination of them significantly reduced 786O cell viability (**Fig. 6C**). The combination effect of belzutifan and rTRAIL was tested using the Chou-Talalay method^42^, where constant ratio combinations of belzutifan and rTRAIL were used to treat 786O to construct the fraction affected vs. combination index (FA-CI) plot using the CompuSyn software. Based on the principle that CI values less than 1 are considered as having a synergistic effect, we concluded that belzutifan and rTRAIL displayed a synergistic cytotoxic effect across most of the tested concentration range, i.e., belzutifan sensitized 786O cells to killing by rTRAIL (**Fig. 6C**). This result confirms our hypothesis that HIF2α inhibition enhances the effectiveness of rTRAIL in killing 786O cells and has clinical implications in treating ccRCC.

**Figure 6.**
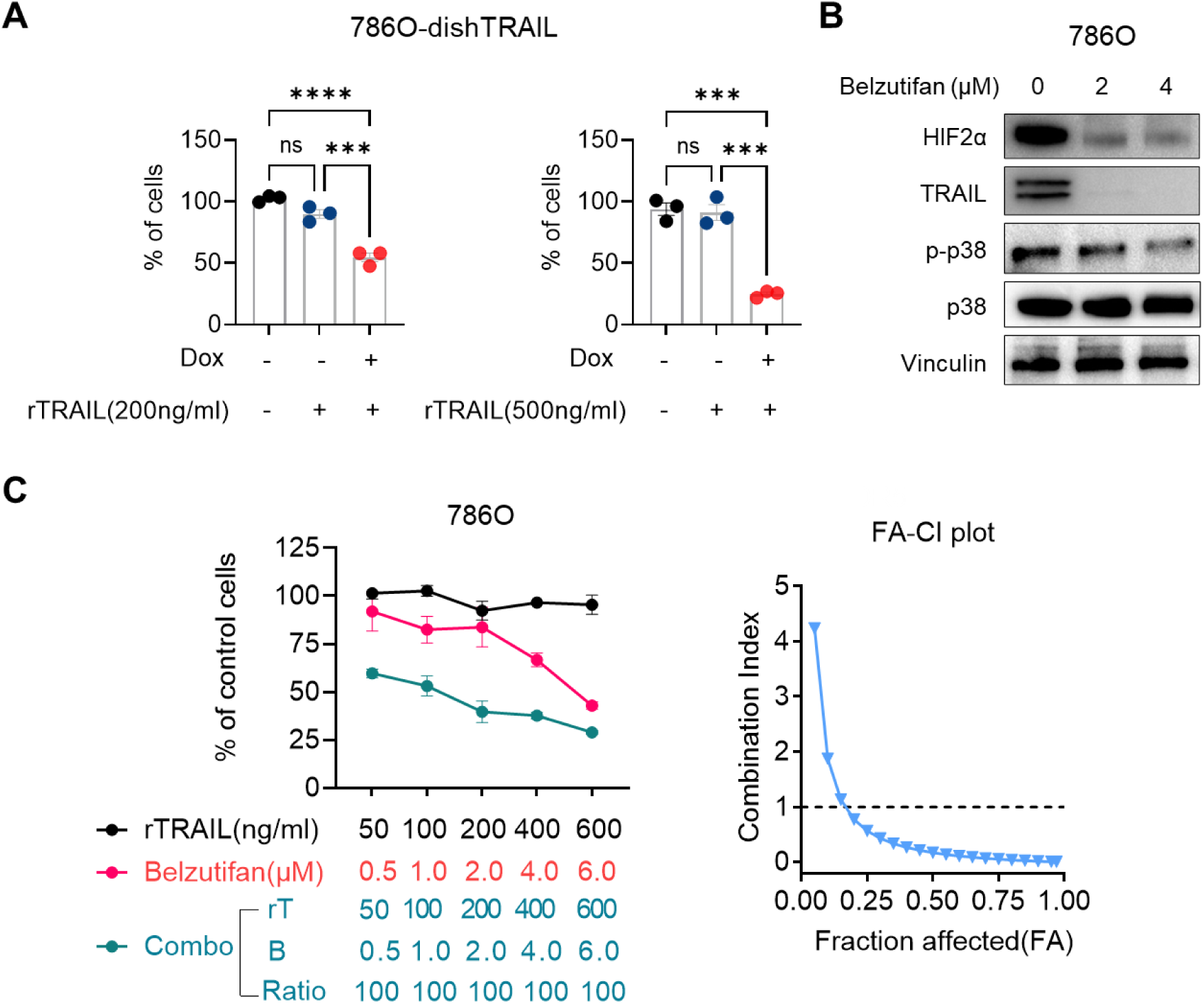
HIF2α inhibitor synergizes with recombinant TRAIL to reduce 786O viability. **A.** Normalized percentages of viable 786O-dishTRAIL cells treated with doxycycline and untreated control (48 hours) followed by treatment with two different concentrations of rTRAIL for 24 hours. Error bars represent SEM. Ordinary one-way ANOVA test with Tukey’s multiple comparison correction, ***P < 0.001, ****P < 0.0001. **B.** Immunoblotting of HIF2α, TRAIL, p-p38, and p38 in 786O treated with 2μM or 4μM belzutifan for 48 hours. **C.** Cell viability quantification of 786O under three treatments: incremental doses of rTRAIL (black), incremental doses of belzutifan (red), and combinations of rTRAIL and belzutifan at a fix ratio of 100 (cyan). The results were analyzed with CompuSyn software to construct FA-CI plot (right). CI<1 indicates synergism effect.

## Discussion

Our investigation into the synthetic essentiality of TRAIL in VHL-deficient ccRCC uncovers critical insights into the molecular interplay between TRAIL and HIF2α signaling pathways. Our results demonstrate that TRAIL acts as a direct transcriptional target of HIF2α and plays a paradoxical role in promoting ccRCC cell proliferation, primarily through the activation of the p38 MAPK pathway and facilitation of the G1/S transition. This pro-tumorigenic role of TRAIL is further supported by its requirement for sustaining HIF2α expression and activity, revealing a reciprocal regulatory loop. This counterintuitive role of TRAIL pathway was initially hinted by the pathological findings that high expressions of TRAIL and DR5 were independent prognostic factors of poor clinical outcomes in patients with ccRCC ^43^. Both cancer-cell-intrinsic (pro-proliferation) and extrinsic mechanisms (cytokine production and immunosuppression) were speculated to explain how TRAIL signaling may promote ccRCC^16^. Our study with human ccRCC cells provides compelling evidence for the cancer cell autonomous role of TRAIL signaling (although non-autonomous roles may also contribute).

Previous studies have identified various targets for VHL-deficient ccRCC, yet our findings highlight TRAIL as a novel synthetic essential gene, opening new avenues for targeted therapy. Besides the VHL-HIF2α axis, oncogenic KRAS also converts TRAIL/TRAIL-R signaling from tumor-restricting to metastasis-promoting in colorectal cancer, lung cancer and pancreatic cancer where KRAS mutation is frequent ^44,45^. KRAS mutation is rare in ccRCC ^46^, and the synthetic essential conversion of TRAIL signaling in ccRCC is through a distinct HIF2α-dependent mechanism, highlighting the versatility of regulatory circuits for TRAIL signaling. Our results with RIPK1 inhibitor support the involvement of RIPK1-incorporated secondary signal complex downstream of TRAIL to mediate p38 phosphorylation and subsequent cell cycle regulation. Considering that RIPK1 is a target of EGLN1–pVHL and can be activated under prolonged hypoxia ^47^, pseudohypoxia in VHL-null ccRCC may contribute to enabling RIPK1 as a downstream effector of TRAIL signaling.

Moreover, the discovery that TRAIL depletion re-localizes DR5 to the plasma membrane offers a novel mechanistic insight into how ccRCC cells modulate TRAIL signaling and evade TRAIL-induced apoptosis. This finding not only elucidates a potential resistance mechanism but also portraits a unique therapeutic scene: targeting endogenous TRAIL signaling enhances the efficacy of exogenous TRAIL-based therapy by increasing the availability of its receptors on the cell surface. Our results echo previous report on TRAIL-dependent nuclear trafficking and chromatin association of TRAIL-R in pancreatic cancer cells ^48^, although nuclear localization of DR5 was not observed in 786O cells. Since the subcellular localization of DR5 functions as a switch to determine the cellular sensitivity to TRAIL and other ligand-independent functions ^40^, manipulation of DR5 localization may be a key aspect to effectuate TRAIL therapy in the future.

The limitations of our study include the use of a single ccRCC cell line 786O (the primary reason is the difficulty in obtaining the other top-ranked ccRCC cell lines that show preferential essentiality for TRAIL, such as TUHR4TKB, KMRC2, SLR24) and the immunodeficient context of the study. Nevertheless, because 786O is the most commonly used and arguably the most robust model for VHL-deficient ccRCC that enabled so many groundbreaking discoveries in ccRCC and hypoxia, we believe that our conclusions are solid and generalizable.

Taken together, our study identifies TRAIL as a synthetic essential gene in VHL-deficient ccRCC, mediated through a novel feedback loop with HIF2α that promotes tumor progression. The observed sensitization of ccRCC cells to rTRAIL cytotoxicity upon TRAIL depletion or HIF2α inhibition suggests a promising strategy for overcoming the therapeutic challenges of TRAIL resistance and belzutifan resistance. The recent effort to make TRAIL therapy work for oncology ^17^ just had its first success ─ approval of Aponermin (a form of rTRAIL) in combination with thalidomide and dexamethasone in China as the third-line treatment of patients with relapsed or refractory multiple myeloma ^49^. Using TRAIL therapy (rTRAIL or TRAIL-R agonists) in a combinatorial setting is most likely also the path for TRAIL therapy to work in solid tumors. In particular, our study illuminates a clinical path for combining TRAIL therapy (possibly Aponermin) with HIF2α inhibitor belzutifan in treating VHL-deficient ccRCC.

## Methods

### Cell culture

786O and 293T were purchased from the American Type Culture Collection (ATCC, CRL-1932, CRL-3216). 786O and its sublines used in this study were cultured in RPMI1640 medium (GE Healthcare, SH30027.01) supplemented with 10% fetal bovine serum (GE Healthcare, SH30396.03) and 100 U/mL of Penicillin–Streptomycin (Caisson Labs, PSL01) at 37 °C in a humidified incubator with 5% CO_2_. 293T were maintained in DMEM, supplemented with 10% fetal bovine serum and 100 U/mL of Penicillin–Streptomycin at 37 °C in a humidified incubator with 5% CO_2_. All cells were tested with the Mycoplasma Assay Kit (Agilent Technologies, 302109) to ensure the mycoplasma-free status.

### Animal experiments

All animal works performed in this study were approved by the Institutional Animal Care and Use Committee at the University of Notre Dame. NCr nude female or male mice (6-8 weeks old) were purchased from Taconic Biosciences (Rensselaer, NY). For orthotopic modeling, 2 × 10^6^ viable cells were injected near the lower pole into the renal parenchyma. After tumor transplantation, one group of mice was on a doxycycline diet. Tumors were monitored weekly by bioluminescent imaging with Spectral Ami HT Advanced Molecular Imager (Spectral Instruments Imaging, Tucson, AZ). Mice were sacrificed at the indicated endpoint.

### Expression constructs and lentiviral transfection

Three validated lentiviral shRNAs against human *TNFSF10* in the pLKO backbone were purchased from Sigma Mission with designated target sequences TRCN0000005924 (CCTCAAAGTGACTATTCAGTT), TRCN0000005925 (GCTGTAACTTACGTGTACTTT), and TRCN0000005927 (CCATTTCTACAGTTCAAGAAA). The control shRNA (SHC202) was also acquired from Sigma. For inducible TRAIL silencing, the *TNFSF10*-targeted shRNA with the sequence GCTGTAACTTACGTGTACTTT was synthesized using the following primers: Forward primer:

5’-CCGGGCTGTAACTTACGTGTACTTTCTCGAGAAAGTACACGTAAGTTACAGCTTTTT-3’,

Reverse primer:

5’-AATTAAAAAGCTGTAACTTACGTGTACTTTCTCGAGAAAGTACACGTAAGTTACAGC-3’.

These oligos were cloned into the AgeI (New England Biolabs, #R0552S) and EcoRI (New England Biolabs, #R0101S) digested Tet-pLKO-puro (Addgene #21915) . For CRISR-Cas9 knockout of HIF2α, we sgRNA with the following primers ^10^:

sgHIF2α forward primer:5’-CACCGTCATGAGGATGAAGTGCA-3’, reverse primer: 5’-AAACTGCACTTCATCCTCATGAC-3’, sgControl forward primer: 5’-CACCGCTTGTTGCGTATACGAGACT-3’, reverse primer: 5’-AAACAGTCTCGTATACGCAACAAG-3’. The sgRNA was cloned into the BsmBI (New England Biolabs, #R0739S) digested pLenti-CRISPRv2 puro vector (Addgene, #39481).

Parental cells were infected with lentivirus packaged by 293T cells. Stable inducible shRNA knockdown cells or sgHIF2α knockout cells were selected with puromycin (4μg/mL) followed by expression verification.

### ChIP-seq data re-analysis

ChIP-seq raw data ^27^ was retrieved from Gene expression Omnibus under the accession code GSE98015. Sequencing reads were aligned to hg19 using bowtie2, and samtools was used to convert sam files to sorted bam files. HIF2α peaks were called using MACS2 with options *– broad* and -*p 1e-8* and the corresponding input samples as controls. Normalized bigwig files called peaks were loaded to IGV for visualization.

### Chromatin Immunoprecipitation (ChIP) followed by qPCR

For ChIP-qPCR, samples were collected following Rodrigues et al. ^27^. Briefly, cells were cross-linked for 10 minutes at room temperature in 1% formaldehyde in growth media, followed by 5 minutes of quenching with 0.125 mol/L glycine. Cells were washed twice with PBS and lysed using the lysis buffer (20 mmol/L Tris–HCl pH 8.0, 150 mmol/L NaCl, 2 mmol/L EDTA pH 8.0, 0.1% SDS, and 1% Triton X-100). Samples were then incubated overnight with the following antibodies at 4°C: HIF2α (Cell Signaling Technology, #59973) and control rabbit polyclonal IgG (Cell Signaling Technology, #2729). 30μL Protein G Sepharose 4 Fast Flow beads (Sigma, GE17-0618-01) were then incubated with the sample slurry at 4°C for 2 hours. DNA was eluted in elution buffer (50 mmol/L NaHCO3, 1% SDS), and cross-links were reversed for 1 hour at 65°C. DNA was purified using the DNA purification spin columns (Cell Signaling Technology, 14209). The samples were sonicated in a Covaris S220 Ultrasonicator System for 12 minutes to obtain a fragment size between 150–300bp. To validate individual binding, qPCR was performed using SYBR Green qPCR master mix (Bimake, B21202). The enrichment of the *TNFSF10* and *SLC2A1* promoter regions was calculated relative to the IgG control. For the *TNFSF10* promoter region, forward primer 5’-AACACGGGAGAAGACCAA-3’; reverse primer 5’-TTCCTGCACGTTTCCCATTC -3’. For the *SLC2A1* promoter region, forward primer 5’-TTTCCACCTGCTGCAATCCT -3’; reverse primer 5’-ATCCTGAGCCTCCCTCTCTG -3’.

### RNA-seq and data analysis

RNA was extracted in duplicates from 786O-Ctrl, 786O-sgEPAS, and 786OdishTRAIL with and without doxycycline cells using the Quick-RNA Miniprep Kit (Zymo Research, #R1054) following the manufacturer’s manual. RNA quality was assessed with Agilent Bioanalyzer 2100. Purified RNA (1μg) was used for the Illumina NovaSeq 6000 Sequencing (Novagene). RNA-seq sequencing reads were aligned to hg19 using STAR aligner, and the DESeq2 package was used to calculate differential expression genes. Gene Set Enrichment Analysis (GSEA) was conducted using the normalized read counts from each sample to assess enrichment across the Human MSigDB Hallmark gene sets. Pathways identified as significant (FDR < 0.25) were selected for further visualization using R package.

### TCGA, CCLE, and CPTAC data analysis

The TCGA RNA-seq transcriptomic data for kidney renal cell carcinoma (KIRC) were downloaded from cBioportal. The transcriptomic data for the Cancer Cell Line Encyclopedia (CCLE) were downloaded from DepMap portal. Gene expressions were extracted using Expression public 23Q4 datasets. Expression of the genes of interest was normalized and log2 transformed. Hypoxia scores were computed as the geometric mean signature expression of the MSigDB human Hallmark hypoxia gene set. Protein expression of ccRCC cohorts was downloaded from the Clinical Proteomic Tumor Analysis Consortium (CPTAC) portal. Data was normalized prior to correlation analysis. The correlation analyses in the study were conducted using either the Pearson or Spearman methods, considering the specific instances.

### Quantitative RT-PCR

Total RNA was extracted from cells using the Total RNA Isolation Miniprep kit (BioBasic, #BS1361) following the manufacturer’s instructions. Total RNA was used for reverse transcription to synthesize cDNA by using All-in-One cDNA Synthesis SuperMix (Bimake, #B24403). qPCR was then performed using SYBR Green qPCR master mix (Bimake, B21202) with the following primers: VEGFA forward primer: 5’-AGGGCAGAATCATCACGAAGT-3’, reverse primer: 5’-AGGGTCTCGATTGGATGGCA-3’; EPAS1 forward primer: 5’-GTCGACCTCATACTTCTCGT-3’, reverse primer: 5’-CAAAGGGCAGCTCCCACCCCTGAG-3’; SLC2A1 forward primer: 5’-ATTGGCTCCGGTATCGTCAAC-3’, reverse primer: 5’-GCTCAGATAGGACATCCAGGG-3’; 18s forward primer: 5’-GGAGTATGGTTGCAAAGCTGA-3’, reverse primer: 5’-ATCTGTCAATCCTGTCCGTGT-3’; RPL13A forward primer: 5’-CATAGGAAGCTGGGAGCAAG-3’, reverse primer: 5’-GCCCTCCAATCAGTCTTCTG-3’. Relative expression levels were calculated based on the 2^−ΔΔCT^ method.

### Colony formation and cell proliferation assays

For the 2D colony formation assay, 1,500 cells per well were seeded into 6-well plates and incubated for 7∼14 days, with the medium refreshed every 2-3 days. For 12-well plates, 1,000 cells per well were seeded. Cells were fixed and stained with 0.2% crystal violet/25% methanol solution for 1 hour at room temperature, and the colony number with diameters of colony > 0.2 cm per well was counted. In the cell proliferation assay, cell growth was monitored using a liver cell imaging system (LS720, Etaluma). Images were captured at 4-hour intervals. Cell confluency was quantified using the Lumaquant 8.8 Image analysis software.

### MTT assay and drug synergism analysis

For MTT (3-(4,5-dimethylthiazol-2-yl)-2,5-diphenyltetrazolium bromide) assays, cells (1000/well) were seeded in 96-well plates under triplicates or quadruplicates, and treated with various concentrations of a single treatment of HIF2α inhibitor belzutifan (Ambeed Inc., #A1216748), recombinant human TRAIL (BioLegend, #752902), or combinational treatment of two agents for designated durations (24 hours to 48 hours). Cells were incubated with MTT for 2 to 4 hours, and then formazan crystals were re-solved in Dimethyl sulfoxide (DMSO, Sigma-Aldrich, #D4540). The 96-well plate was read at 570 nm absorbance by a microplate reader (Epoch2, Biotek). The data for drug synergism analysis were generated by the MTT assay. Cells were plated a day prior to being subjected to both individual and combined drug treatments, with each drug concentration (spanning five serial dilutions) replicated three times. Following this, the absorbance of each well was adjusted relative to control wells treated with DMSO to determine the drug’s impact, quantified as Fraction Affected (FA) by the treatment. CompuSyn software is used to generate a simulation for the drug synergism analysis. The CI was plotted along with the indicated FA for each cell line. CI > 1, antagonism; CI = 1, addition; CI < 1, synergism.

### Flow cytometry

Cell cycle analysis was done by genomic DNA staining using propidium iodide (ThermoFisher Scientific, P1304MP). Cells were treated with 100 nM nocodazole for 16 hours to synchronize cells prior to doxycycline induction. Cells were collected and fixed with the 70% ethanol at 4 °C for 30 minutes. After washing with PBS (Cytiva, #SH30028) three times, cells were stained with propidium iodide at room temperature for 30 minutes. For the apoptosis assay, cells were stained with APC-conjugated Annexin V (Cytek Biosciences, #20-6409-T100) and DAPI (Sigma-Aldrich, #D9542) in 1X Annexin V binding buffer according to manufacturer instructions (R&D Systems, 4830-01-K). Cells treated with 1 µM staurosporine for 4 h were used as the positive control for apoptosis. For the detection of the expression of cell surface DR5, cells were stained with anti-DR5 (Biolegend, #307302) followed by secondary antibody conjugated with the Alexa Flour 647 (Jackson ImmunoResearch Laboratory, 115-605-146). Fluorescence was detected and quantified on Beckman Coutler CytoFLEX S flow cytometer. All FACS files were analyzed using FlowJo software.

### Immunofluorescence assay

Cells were seeded on a coverslip in the 24-well plate and cultured till 70% confluency was achieved. Cells were washed and fixed with 4% formaldehyde for 15 minutes at room temperature. After fixation, cells were permeabilized and blocked using a blocking buffer (3% BSA + 0.3% Triton X-100 in PBS) for 1 hour. Cells were incubated with the following primary antibodies: TRAIL (Cell Signaling Technology, #3219), Lamp1 conjugated with Alexa Fluor647 (Santa Cruz, SC-19992), and DR5 (Biolegend, #307302) at 4 C for 16 hours. Next, the cells were stained with secondary antibodies Alexa Fluor 647 anti-mouse (Jackson ImmunoResearch Laboratory, 115-605-146) and Alexa Fluor 488 anti-rabbit (Life Tech, #A11008) at 1:500 dilution for 1 hour at room temperature. The nucleic was stained using 1 µg/ml DAPI (Sigma-Aldrich, D9542). Cells were mounted on glass slides using mounting media (ThermoFisher Scientific, SP15100) before fluorescence confocal microscopy using Nikon A1R HD Confocal Microscope.

### Western blotting

The western blot procedure was conducted as we described ^50^. Briefly, cells were lysed in RIPA buffer containing protease inhibitors and phosphatase inhibitors. All samples were run with standard SDS-PAGE. Primary antibodies included: TRAIL (Cell Signaling Technology, #3219), HIF2α (Cell Signaling Technology, #59973), p38 (Cell Signaling Technology, #9212), Phospho-p38 MAPK (Thr180/Tyr182) (Cell Signaling Technology, #4511), RB (Cell Signaling Technology, #9309), Phospho-RB (Ser780)(Cell Signaling Technology, #8180), cyclin D1(Cell Signaling Technology,#2978P), cyclin D3 (Cell Signaling Technology, #2936), cyclin E1 (Cell Signaling Technology,#4219), E2F1 (Novus Biologicals, #NBP2-37474), Phospho-IκBα (Cell Signaling Technology, #2859), IκBα (Cell Signaling Technology, #4814), phospho-NF-κB p65 (Ser536) (Cell Signaling Technology, #3033), p-ERK (Cell Signaling Technology, #4370P), ERK (Cell Signaling Technology, #4695). β-actin (Santa Cruz, #sc-47778), Vinculin (Sigma-Aldrich, #05-386). Secondary antibodies included HRP-linked anti-rabbit IgG (Cell Signaling Technology, #7074) and HRP-linked anti-mouse IgG (Cell Signaling Technology, #7076).

### Inhibitor treatment

Cells at 10^5^ per well were plated in a 6-well plate and allowed to grow until they reached 60∼80% confluency. At this stage, the cells were treated with Necrostatin-1s (Cell Signaling Technology, #17802) at 10μM for 1 hour or the p38 inhibitor SB203580 (ApexBio, #A8254) at 25μM for 1h or 6h. Cells were then harvested for western blot.

### Statistical analysis

Experiments were performed two or three times, and conclusions were drawn only when the results were reproducible. One representative result among these replicates is shown in the figures. For non-omics data, statistical analyses were performed using GraphPad Prism v9.3 (RRID: SCR_002798). All data are presented as mean ± SEM (standard error of the mean). We followed this workflow for statistical testing: Shapiro-Wilk test was performed to assess for normality of data distribution: (i) in case of normality, when only two conditions were to test, we performed unpaired t-test; when more than two conditions were to compare, we performed a parametric one-way or two-way ANOVA followed by post hoc test with recommended correction for multiple comparisons to assess the significance among pairs of conditions. (ii) in case of non-normality, when only two conditions were to test, we performed a Mann-Whitney U test; when more than two conditions were to compare, we performed a non-parametric one-way ANOVA followed by a recommended test to assess the significance among pairs of conditions. Sample sizes, error bars, P values, and statistical methods are noted in the figures or figure legends. Statistical significance was defined as P < 0.05 unless otherwise noted.

## Data availability statement

Raw RNA-seq data are deposited in the Gene Expression Omnibus (GEO) (RRID: SCR_005012) under accession code GSE259401. All other raw data and the materials generated in this study are available upon request to the corresponding author.

## Acknowledgments

We thank the Lu lab members for their essential comments and suggestions during this work. We are grateful for the support from core facilities used in this study. This work was supported by a research grant from VHL Alliance’s Competitive Research Grant Program (Xin Lu), a grant from Naughton Faculty Research Accelerator Program at University of Notre Dame (Xin Lu, E. Szegezdi), and Boler Family Foundation (Xin Lu). Other support included National Institutes of Health grants R01CA248033 and R01CA280097 (Xin Lu), Department of Defense grants W81XWH2010312, W81XWH2010332, HT94252310010 and HT94252310613 (Xin Lu), and National Institutes of Health grant R01CA262439 (Z. Schafer).

## Author Contributions

**X. Wang:** Conceptualization, investigation, methodology, data curation, formal analysis, validation, visualization, manuscript writing. **L. Duong, Y. Qin, R. Parrotta, P.K. Purohit:** Investigation, methodology, data curation, formal analysis, validation, visualization. **Y. Fang, G. Liu, J. Wen, Y. Liu, Y. Zhang, J. Zhao:** Investigation and methodology. **J. He, Z.T. Schafer:** Methodology. **Xuemin Lu, E. Szegezdi:** Supervision. **Xin Lu:** Conceptualization, investigation, formal analysis, resources, project administration, supervision, funding acquisition, manuscript writing.

