## Supplemental Figures for "Synthetic essentiality of TRAIL/TNFSF10 in VHL-deficient renal cell carcinoma"

### Supplementary Figures

#### Supplementary Figure S1

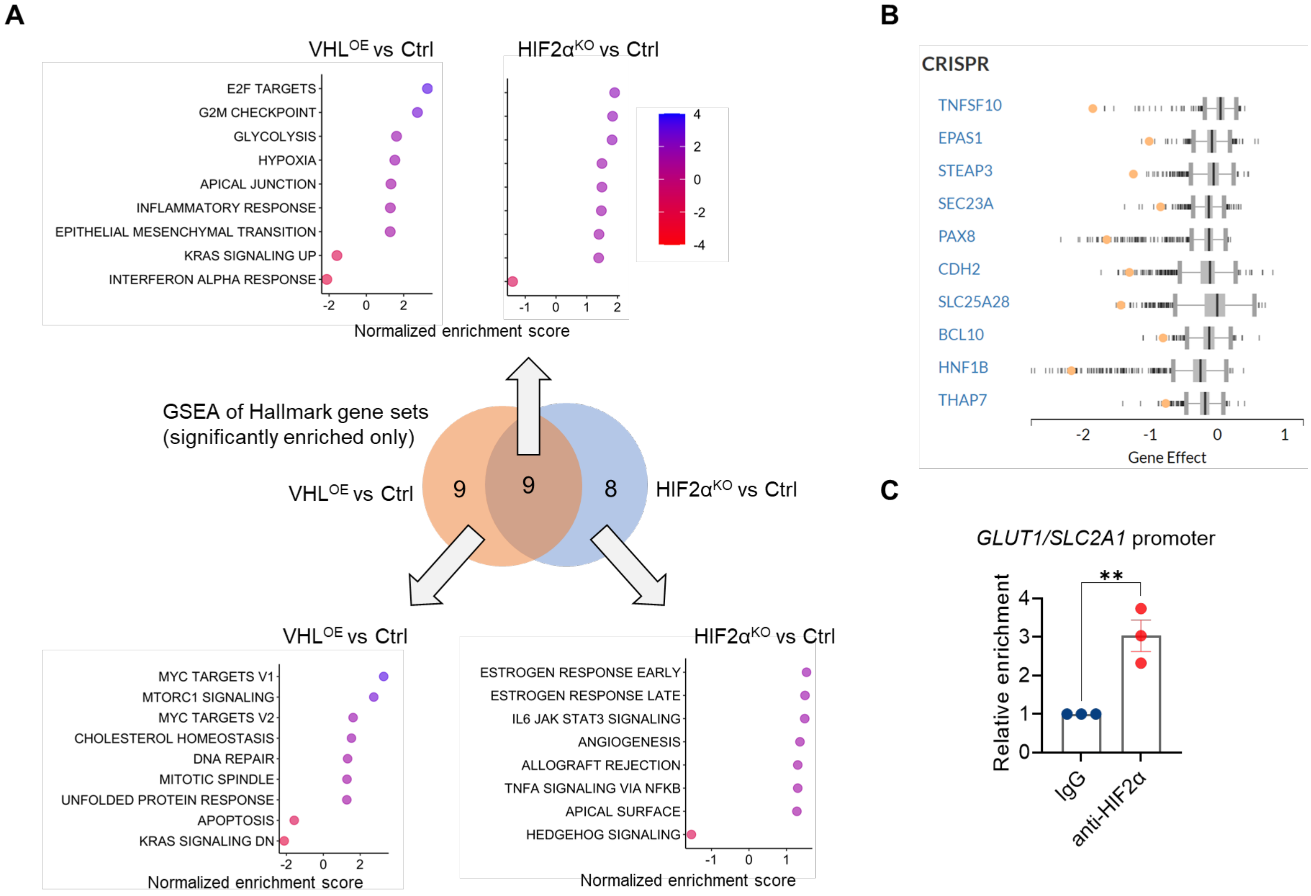


**Supplementary Figure S1. TRAIL is a HIF2α target and a candidate synthetic-essential.** **A**. Gene Set Enrichment Analysis (GSEA) result showing MSigDB hallmark pathways (P < 0.05, FDR < 0.25) enriched for the differentially expressed genes between two sets of conditions: (1) 786O-sgEPAS1 (HIF2α knockout) and 786O-Ctrl (HIF2α$^{KO}$ vs. Ctrl) and (2) 786O-VHL overexpression and 786O-Ctrl (VHL$^{OE}$ vs. Ctrl), segregated by Venn diagram to three plots. The color of the bubbles corresponds to the FDR. **B**. Box plot showing gene effect scores of top 10 preferential essential genes in 786O cell line, generated based on CRISPR screening by the cancer dependency map (DepMap) project. The gene effect score for each gene in 786O is indicated by the yellow dot. The vertical lines represent the distribution of gene effect scores across other cell lines for each gene. **C**. ChIP-qPCR result comparing the association of IgG and HIF2α to the *GLUT1/SLC2A1* promoter in 786O (n=3). Error bars stand for SEM, unpaired t-test, **P < 0.01.

### Supplementary Figure S2

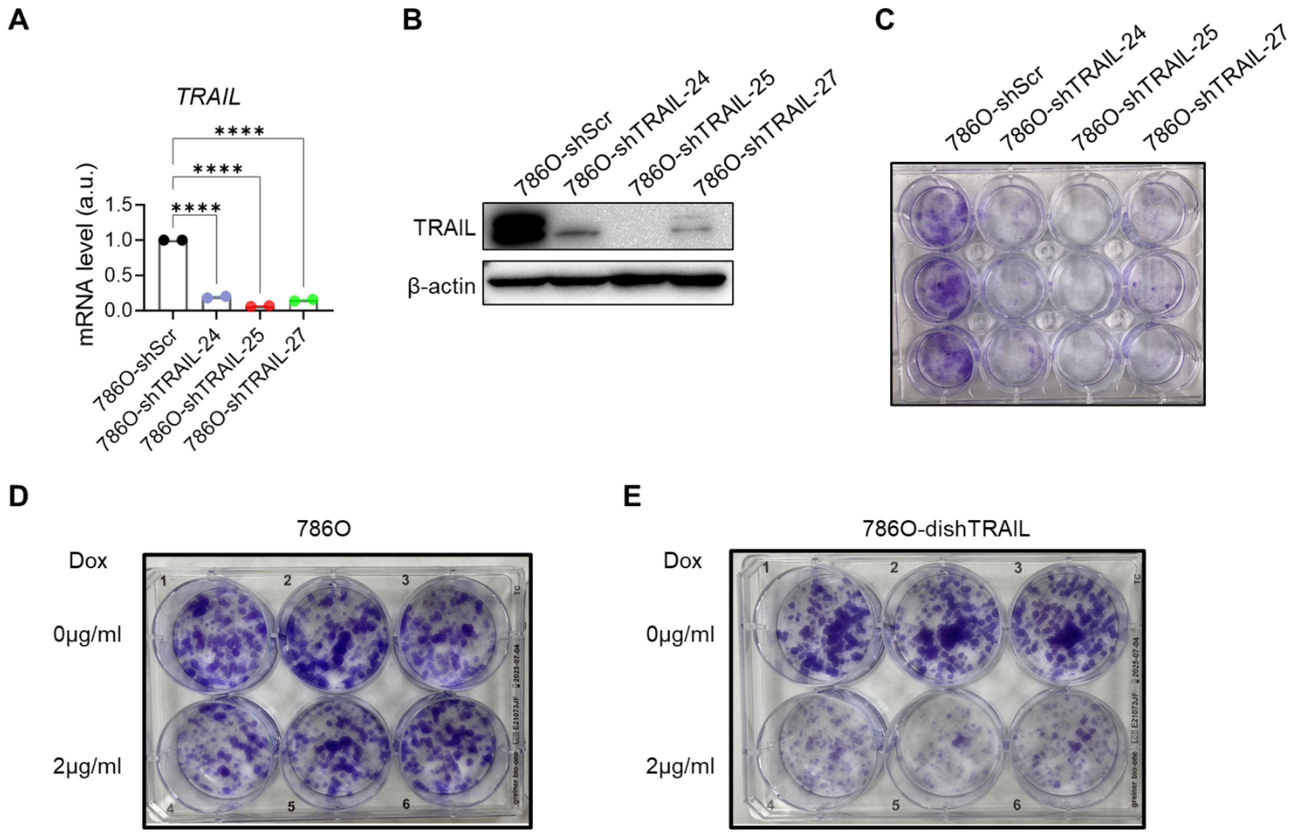


**Supplementary Figure S2. TRAIL depletion attenuates ccRCC proliferation in vitro.**

**A.** qRT-PCR result showing *TNFSF10* mRNA levels for 786O cells stably transduced with shRNA targeting scramble (control) or TRAIL (#24, #25, #27). Unpaired t-test, ****P < 0.0001. **B**. Western blot of TRAIL for 786O cells stably transduced with shRNA targeting scramble (control) or TRAIL (#24, #25, #27). **C**. Clonogenic assay for 786O cells stably transduced with shRNA targeting scramble (control) or TRAIL (#24, #25, #27). **D-E**. Clonogenic assay for 786O or 786O-dishTRAIL cells under doxycycline off or on (2 μg/ml, 10 days) conditions.

### Supplementary Figure S3

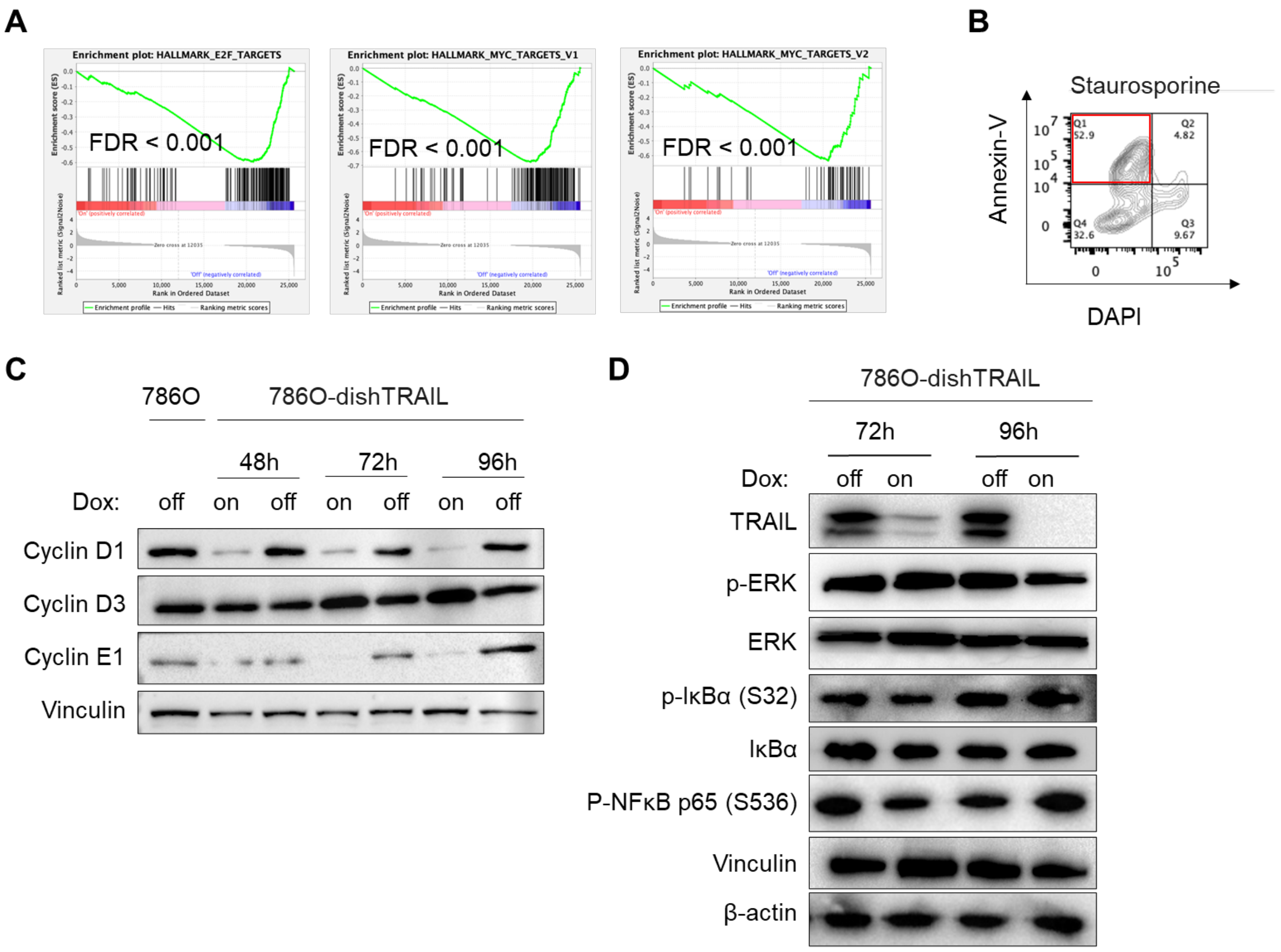


**Supplementary Figure S3. TRAIL activates p38 MAPK and facilitates G1/S progression in ccRCC cells.**

**A.** GSEA enrichment plots of top enriched MSigDB hallmark pathways. **B.** Representative flow cytometry plots of the apoptosis assay using annexin-V-APC and DAPI staining on 786O cells treated with staurosporine (1 μM, 4 hours) as positive control. The apoptotic cell population is indicated in the red box. **C**. Immunoblotting of cyclin D1, cyclin D3, and cyclin E1 for 786O and 786O-dishTRAIL treated with doxycycline (Dox on) and untreated controls (Dox off) across three time points (48, 72 and 96 hours). **D**. Immunoblotting of TRAIL, p-ERK, ERK, p-iκB-α(S32), iκB-α, p-NF-κB p65 (Ser536) in 786O-dishTRAIL treated with doxycycline (Dox on) and untreated controls (Dox off) across two time points (72 and 96 hours).

**Supplementary Figure S4**

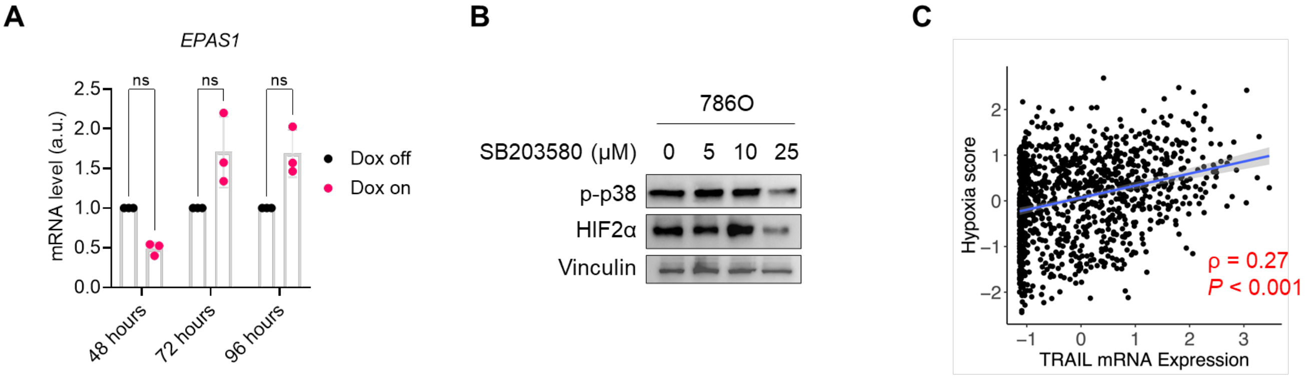


**Supplementary Figure S4. TRAIL is required to sustain HIF2α expression and activity in ccRCC cells.**

**A.** qRT-PCR quantification of *EPAS1* mRNA expression in 786O-dishTRAIL treated with doxycycline (Dox on) and untreated controls (Dox off) across three different time points: 48,72 and 96 hours. ns = not significant, two-way ANOVA test with Sidak's multiple comparison correct. **B**. Immunoblotting of phosphorylation of p38 and HIF2α protein expression in 786O treated with 0, 5, 10 and 25μM SB203580 for 1 hour. **C**. Scatter plot of normalized TRAIL expression and Hypoxia scores in cell lines based on the CCLE database. Spearman correlation coefficient ρ and P value are indicated.

### Supplementary Figure S5

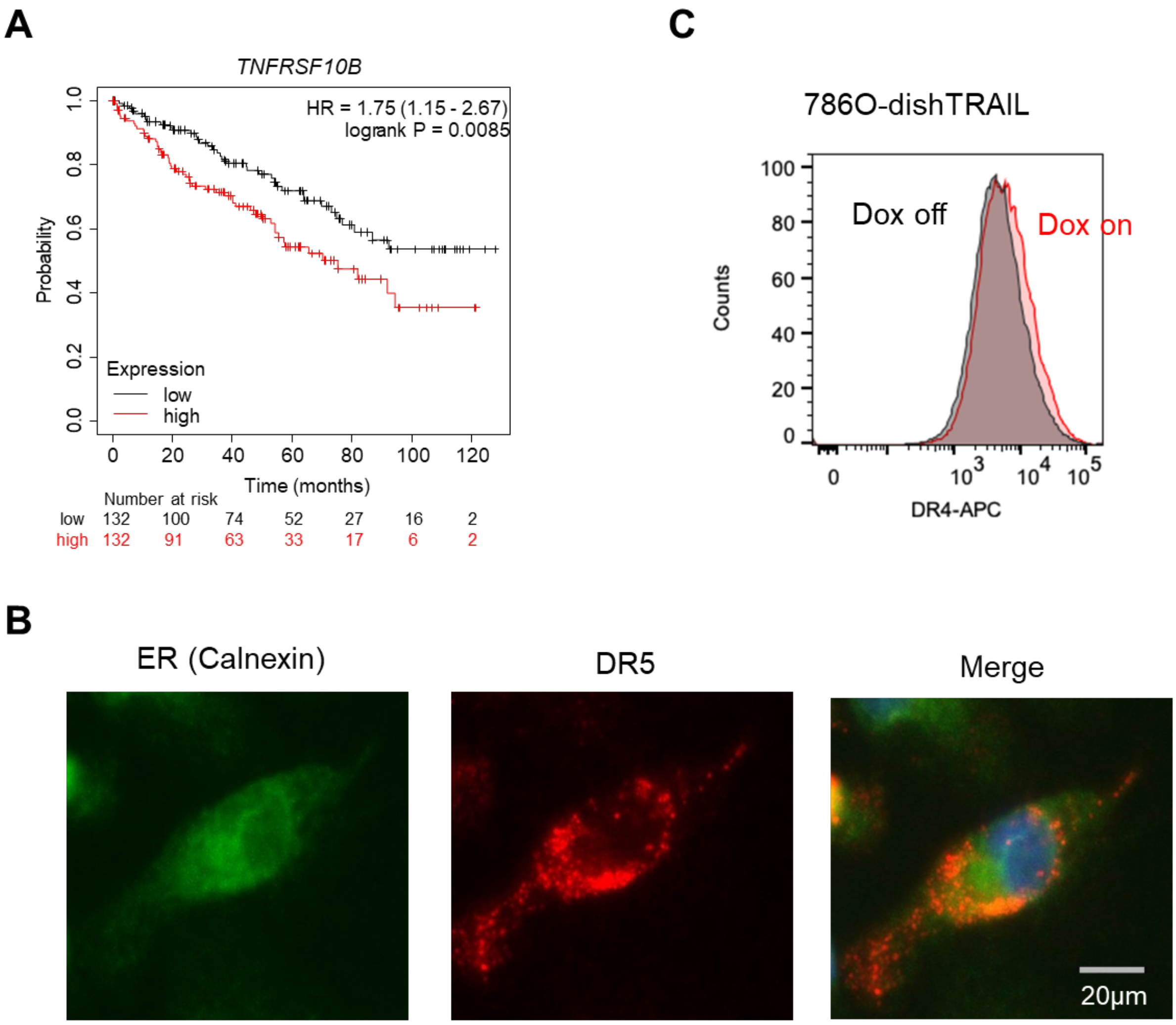


**Supplementary Figure S5. TRAIL depletion re-localizes DR5 to the plasma membrane. A.** Association of *DR5 (TNFRSF10B)* expression levels with overall survival (OS) in TCGA KIRC cohort, drawn with KMplot. **B**. Immunofluorescence co-staining of ER marker calnexin and DR5 in 786O cells. Scale bar 20μm. **C.** Flow cytometry of DR4 expression on the cell surface in 786O-dishTRAIL cells under Dox on or Dox off conditions (48 hours).
